# GONST2 transports GDP-Mannose for sphingolipid glycosylation in the Golgi apparatus of Arabidopsis

**DOI:** 10.1101/346775

**Authors:** Beibei Jing, Toshiki Ishikawa, Nicole Soltis, Noriko Inada, Yan Liang, Gosia Murawska, Fekadu Andeberhan, Ramana Pidatala, Xiaolan Yu, Edward Baidoo, Maki Kawai-Yamada, Dominique Loque, Daniel J. Kliebenstein, Paul Dupree, Jenny C. Mortimer

**Affiliations:** Joint BioEnergy Institute, Emeryville, CA, 94608, USA; BioSciences Division, Lawrence Berkeley National Laboratory, Berkeley, CA 94720, USA.; Graduate School of Science and Engineering, Saitama University, Japan.; Plant Sciences Department, UC Davis, Davis, CA 95616, USA; Graduate School of Biological Sciences, NAIST, Nara, 6300192, Japan.; Department of Biochemistry, University of Cambridge, Cambridge, UK

## Abstract

The Golgi lumen is the site of many different glycosylation events, including cell wall polysaccharide biosynthesis and lipid glycosylation. Transporters are necessary for the import of the substrates required for glycosylation (nucleotide sugars) from the cytosol where they are synthesized. Plants use four GDP-linked sugars to glycosylate macromolecules: GDP-L-Fucose, GDP-D-Mannose, GDP-L-Galactose and GDP-D-Glucose. Of the predicted fifty-one members of the nucleotide sugar transporter/triose phosphate transporter family in Arabidopsis, only four appear to contain the conserved motif needed for the transport of GDP-linked sugars, GOLGI LOCALIZED NUCLEOTIDE SUGAR TRANSPORTER (GONST) 1-4. Previously, we have demonstrated that GONST1 provides GDP-D-Mannose for glycosylation of a class of sphingolipids, the glycosylinositolphosphorylceramides (GIPCs). Here, we characterize its closest homologue, GONST2, and conclude that it also specifically provides substrate for GIPC glycosylation. Expression of *GONST2* driven by the *GONST1* promoter is able to rescue the severe growth phenotype of *gonst1*. Loss of GONST2 exacerbates the *gonst1* constitutive hypersensitive response, as well as the reduced cell wall cellulose content. The *gonst2* mutant grows normally under standard conditions, but has enhanced resistance to the powdery mildew-causing fungus *Golovinomyces orontii*.

## Introduction

The Golgi apparatus is the site of many different glycosylation processes, including the synthesis of polysaccharides, glycoproteins, proteoglycans, and glycolipids. Nucleotide sugars are the universal sugar donors for these processes. In plants, the majority of nucleotide sugars are UDP-linked, but the GDP-linked sugars GDP-D-Mannose (GDP-Man), GDP-D-Glucose (GDP-Glc), GDP-L-Fucose (GDP-Fuc), and GDP-L-Galactose (GDP-Gal) are also critical (Bar-Peled and O’Neill, 2011). Most nucleotide sugars required in the Golgi, including all of the GDP-sugars, are synthesized in the cytosol and therefore need to be translocated into the Golgi lumen via nucleotide sugar transporters (NSTs). To date, some functionally characterized plant NSTs have been reported, (Baldwin et al., 2001; Norambuena et al., 2002; Handford et al., 2004; Bakker et al., 2005; Norambuena et al., 2005; Rollwitz et al., 2006; Reyes et al., 2010; Mortimer et al., 2013; Rautengarten et al., 2014; Rautengarten et al., 2016). *Arabidopsis thaliana* (Arabidopsis) NSTs belong to the NST/triose phosphate translocator (TPT) superfamily which has 51 members that are distributed in six clades (Rautengarten et al., 2014). From this superfamily, only four members, the GOLGI LOCALIZED NUCLEOTIDE SUGAR TRANSPORTER (GONST) sub-clade, are predicted to transport GDP-sugars due to the presence of the conserved GX[L/V]NK motif (Baldwin et al., 2001; Gao et al., 2001; Handford et al., 2004).

Arabidopsis GONST1 was initially identified based on sequence similarity to *Saccharomyces cerevisiae* Vrg4p and *Leishmania donovani* LPG2 GDP-Man transporters (Baldwin et al., 2001). GONST1 can complement the *Saccharomyces cerevisiae vrg4-2* mutant and was the first biochemically characterized plant NST (Baldwin et al., 2001). GONST1 can transport all four plant GDP-sugars *in vitro* (Mortimer et al., 2013). However, analysis of *gonst1* plants revealed a specific role *in vivo* as a GDP-Man transporter which specifically provides substrate for the synthesis of a class of sphingolipids, the glycosylinositolphosphorylceramides (GIPCs; Supplemental Figure S1) (Mortimer et al., 2013). GONST 2-4 were identified as GONST1 homologues on the basis of their sequence similarity to GONST1 (Handford et al., 2004). GONST4 has now been characterized as the Golgi GDP-Fuc transporter and has therefore been renamed GDP-FUCOSE TRANSPORTER1 (GFT1) (Rautengarten et al., 2016). GONST3 has recently been shown to be responsible for GDP-Gal transport and has been renamed GOLGI GDP-L-GALACTOSE TRANSPORTER1 (Sechet et al., submitted). GONST2 was also able to complement *vrg4-2* (Handford et al., 2004) and able to transport all four GDP-linked sugars *in vitro* (Rautengarten et al., 2016), but its function *in planta*, as well as its specificity, remains unknown

GIPC abundance has been long-underestimated, due to their high polarity and poor yield with classic lipid extraction methods. However, recent advances have shown that GIPCs comprise an estimated 64% of plant sphingolipids and ~25% of the total lipids in Arabidopsis leaf, and therefore are probably the predominant sphingolipid class in the plasma membrane (PM) of plant cells (Markham et al., 2006; Markham and Jaworski, 2007; Bure et al., 2011; Cacas et al., 2013; Markham et al., 2013; Cacas et al., 2016). The ceramide is synthesized in the endoplasmic reticulum (ER), where it is either glucosylated to produce glucosylceramides, or it is then trafficked to the Golgi for GIPC biosynthesis. The first GIPC-specific step is the addition of an inositol phosphate to the ceramide via headgroup exchange with a phospholipid phosphatidylinositol by inositolphosphorylceramide (IPC) synthase (IPCS) (Wang et al., 2008). The IPC core is first glycosylated with a glucuronic acid (GlcA) by the CAZy family 8 glycosyltransferase (GT8) INOSITOL PHOSPHORYLCERAMIDE GLUCURONOSYLTRANSFERASE (IPUT1) (Rennie et al., 2014). The GlcA-IPC core is further glycosylated by various other GTs to form mature GIPCs, although the structure and composition varies depending on tissue type and plant species (Markham and Jaworski, 2007; Bure et al., 2011; Tellier et al., 2014; Cacas et al., 2016; Fang et al., 2016; Ishikawa et al., 2016; Ishikawa et al., 2018). In Arabidopsis vegetative tissues, the dominant GIPC carries a Man on the GlcA-IPC, which is added by GIPC MANNOSYL-TRANSFERASE1 (GMT1), from CAZy family GT64 (Fang et al., 2016), the substrate for which is provided, at least in part, by GONST1, and indeed both *gonst1* and *gmt1* have very similar phenotypes (Mortimer et al., 2013; Fang et al., 2016). However, other glycosylated forms of GIPCs have also been identified. For example, in Arabidopsis seeds and pollen, rice, and tobacco leaves, the major GIPC glycosylation is a GlcN(Ac) linked to the GlcA, which can then be extensively decorated (Carter et al., 1958; Kaul and Lester, 1975; Hsieh et al., 1978; Kaul and Lester, 1978; Hsieh et al., 1981; Bure et al., 2011; Tellier et al., 2014; Luttgeharm et al., 2015; Cacas et al., 2016; Ishikawa et al., 2018).

Here, we characterize the function of GONST2 *in planta*. *gonst2* shows a small reduction in GIPC mannosylation but *gonst1gonst2* plants completely lack mannosylated GIPCs. *gonst2* has a growth phenotype indistinguishable from wild type (WT), but *gonst1gonst2* is more severely stunted than *gonst1*, has an early senescence phenotype, dramatically increased salicylic acid, and a decrease in cell wall cellulose content. *gonst2* has increased resistance to the biotrophic pathogen *Golovinomyces orontii*, but not to the necrotrophic pathogen *Botrytis cinerea*. Expression of *GONST2* under the *GONST1* promoter can rescue the *gonst1* phenotype. These data indicate that GONST2 has a similar function to GONST1 in providing GDP-D-Man for GIPC mannosylation.

## Results

### GONST2 is a close homolog of GONST1

Arabidopsis nucleotide-sugar transporters containing a conserved GDP-binding motif (GX[L/V]NK) were first identified by (Baldwin et al., 2001), and named GONST and consist of a clade of 4 proteins (GONST1-2, GONST3/GGLT1, GONST4/GFT1) (Handford et al., 2004) (Supplemental figure S2). A fifth member (GONST5) is found in a distinct clade from GONST1-4 and lacks the GXLNK (Handford et al., 2004; Rautengarten et al., 2014). GONST2 shares 61% identity with GONST1 at the amino acid level, as compared to only 19% with GFT1 (Supplemental Figure S2), and is expressed at a low level in most tissues (Supplemental Figure S3). A single copy of GONST1/2 was found in gymnosperms, non-vascular plants and algae, but both GONST1 and GONST2 orthologs were identified in monocots and dicots (Figure 1). This suggests that the GONST1/2 gene duplication occurred after the Angiosperm-Gymnosperm split, but prior to the monocot-dicot split.

**Figure 1:**
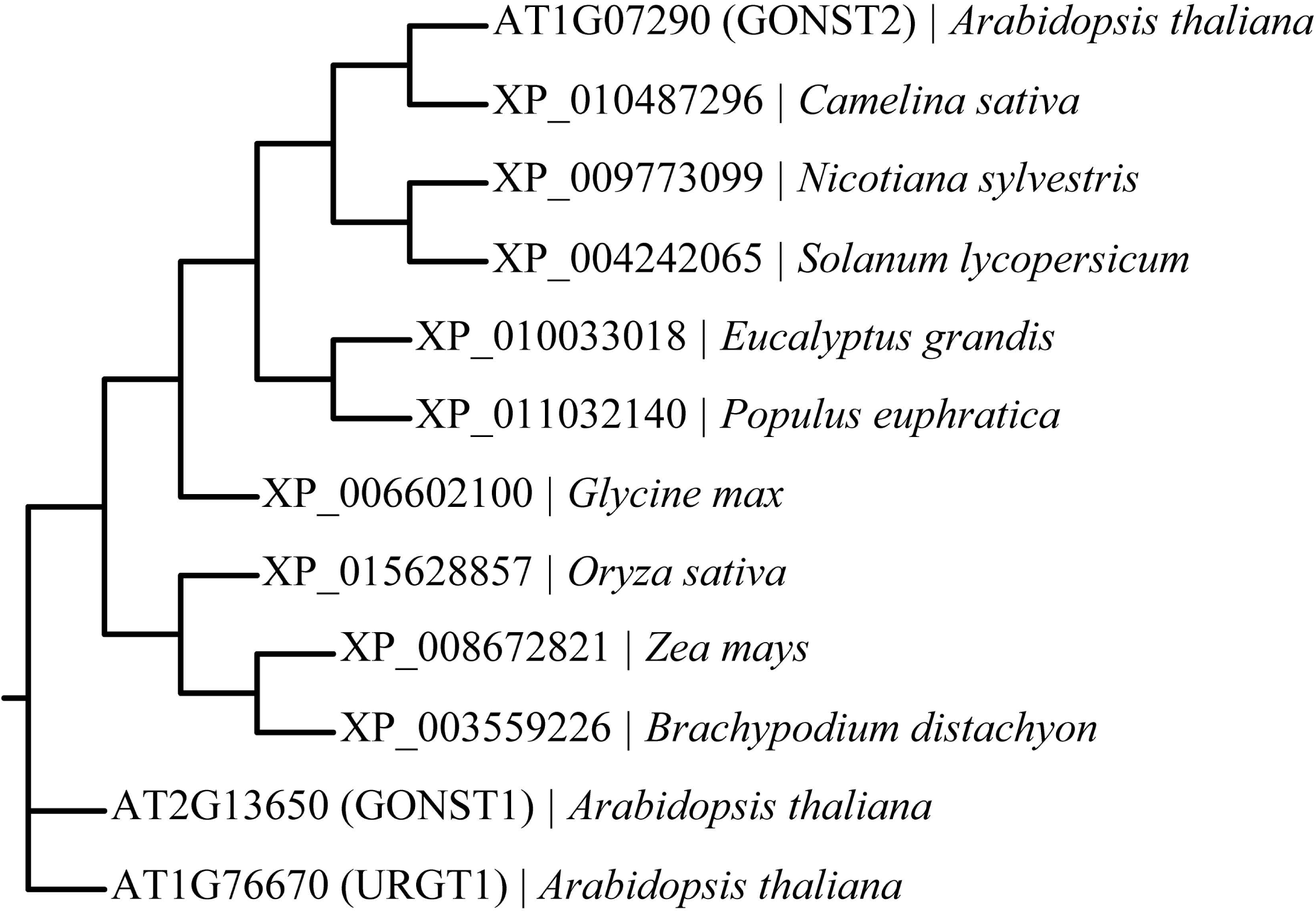
Phylogenetic tree of GONST2 orthologs in plants. The Arabidopsis GONST2 protein sequence was used in a BLASTp search (standard parameters). Sequences were aligned with Clustal Omega (www.ebi.org) (standard parameters). Arabidopsis GONST1 and URGT1 (a UDP-Rha/Gal transporter) were used as outgroups. The Phylip program set (v3.95) was used to build the tree, using standard parameters except where stated, as follows: seqboot (2000 replicates), proml (not rough analysis), consense and drawgram. All bootstrap probabilities were 1.0 with 2000 replicates.

### GONST2 is localized to the Golgi

Previously, attempts to determine the subcellular localization of transiently expressed GONST2 in tobacco under the control of an enhanced cauliflower mosaic virus (CaMV) 35S promoter were unsuccessful (Handford et al., 2004). However, in our hands, using the same promoter, we were able to successfully detect a punctate distribution of transiently expressed GONST2-YFP in tobacco leaves using confocal laser scanning microscopy (Figure 2). To test whether this represented a Golgi-localization, we co-infiltrated the leaves with a cis-Golgi marker, Man49-GFP. Man49-GFP is composed of the first 49 amino acids (aa) of soybean (*Glycine max*) □-1,2 mannosidase I with a C-terminal GFP, which is sufficient for *cis*-Golgi targeting (Saint-Jore-Dupas et al., 2006; Nelson et al., 2007). The majority of the GONST2-YFP signal co-localized with the Man49-GFP signal, suggesting that GONST2 is indeed Golgi localized (Figure 2).

**Figure 2:**
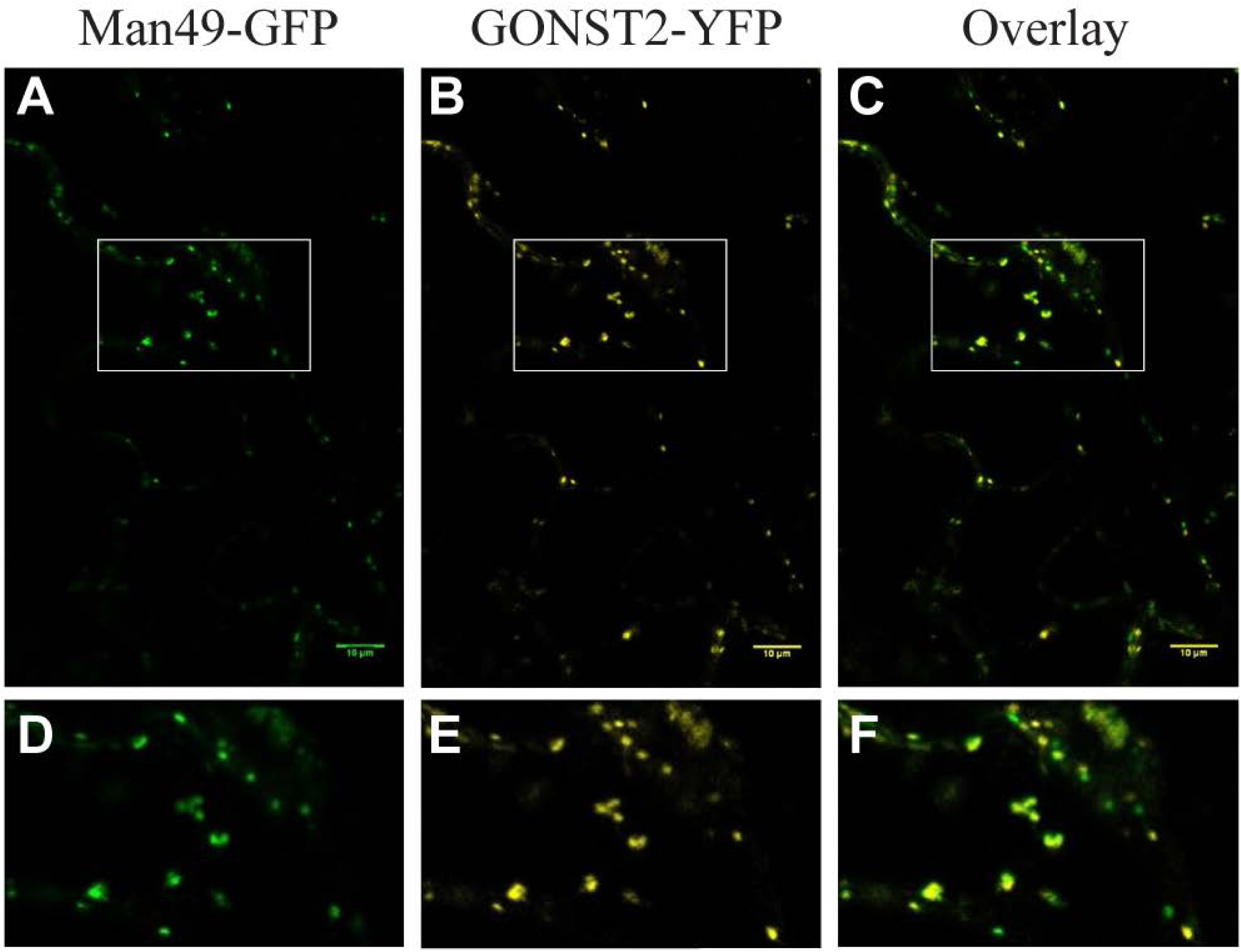
Subcellular localization of GONST2. A *cis*-Golgi marker, Man49-GFP (A, D) along with a C-terminal YFP fusion of GONST2 (B, E), was co-infiltrated into tobacco leaves and expression of the fluorescent fusion proteins detected by confocal microscopy. An overlay of the two channels is shown in (C, F). D, E and F are 63X magnification of the white boxed area in A, B and C. Scale bar = 10 μm.

### Loss of GONST2 exacerbates the *gonst1* growth phenotype

Previously, we isolated and characterized three T-DNA *GONST1* insertion lines (*gonst1-1*, *gonst1-2*, and *gonst1-3*), which all showed a strong developmental phenotype including a dwarf phenotype, poor seed set and spontaneous hypersensitive lesions (SHLs) on leaves (Mortimer et al., 2013). Of these, *gonst1-1* was ecotype Wassilewskija (Ws), and *gonst1-2* and *gonst1-3* were ecotype Columbia-0 (Col-0). We had also isolated a homozygous *gonst2-1* allele (Ws ecotype) which lacked detectable *GONST2* transcript by RT-PCR but did not have a visible phenotype (Figure 3; (Mortimer et al., 2013)). Since no further T-DNA lines were available, here we used CRISPR/Cas9 gene editing to create two further *gonst2* alleles, *gonst2-2* and *gonst2-3*, in the Col-0 ecotype (Supplemental Figure S4). As was the case for *gonst2-1*, *gonst2-2* and *gonst2-3* did not show a visible phenotype compared to WT. However, the *gonst1gonst2* double mutant had a more severe growth phenotype than *gonst1* alone, suggesting redundant function (Figure 3, Supplemental Figure S4, (Mortimer et al., 2013)).

**Figure 3:**
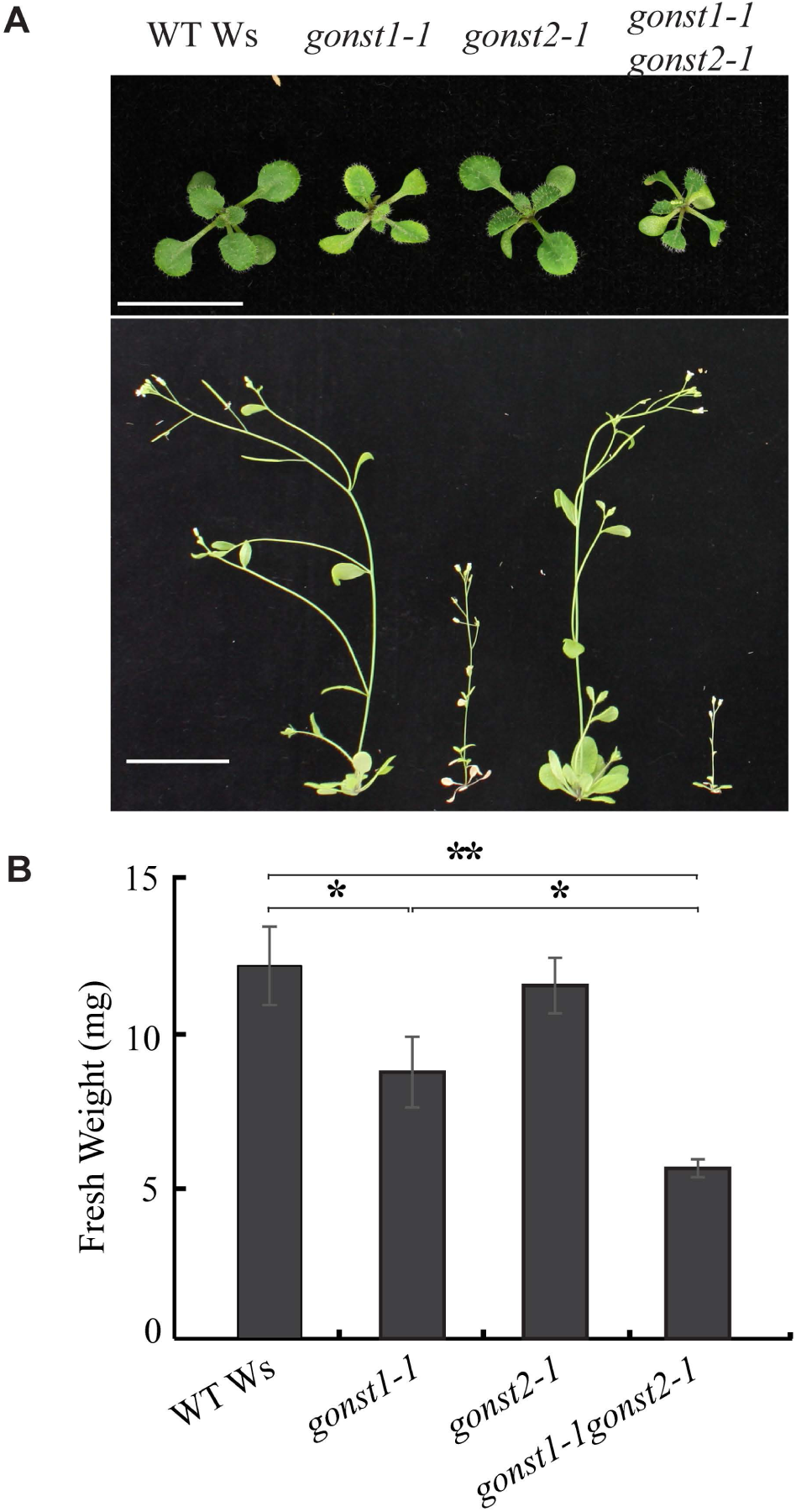
Loss of GONST2 exacerbates the *gonst1* growth phenotype. (A) Top row: 15-day-old, agar grown WT Ws, *gonst1-1*, *gonst2-1* and *gonst1-1gonst2-1* seedlings. Scale bar = 1 cm. Bottom row: 6-week-old WT Ws, *gonst1-1*, *gonst2-1* and *gonst1-1gonst2-1*. Plants were first grown on agar for 10 days, and then transplanted onto soil. Bar = 3 cm. (B) Fresh weight of seedlings of 15-d-old, agar grown WT Ws, *gonst1-1*, *gonst2-1* and *gonst1-1gonst2-1*. *n* = 10 individual plants grown simultaneously, ±SE; asterisk indicate a significant difference from the WT (Student’s *t*-test, * *p* < 0.05, ** *p* < 0.01, *** *p* < 0.001).

### Loss of GONST2 enhances the *gonst1* constitutive hypersensitive response

*gonst1* has biochemical phenotypes consistent with the constitutive activation of plant defense responses, including elevated salicylic acid (SA) and reactive oxygen species (ROS) (Mortimer et al., 2013). To test whether loss of GONST2 also resulted in the constitutive activation of plant defense responses, we measured in situ H_2_O_2_ production (as a proxy for ROS) and SA in *gonst2-1*. *gonst2-1* did not show a significant change in either SA or H_2_O_2_ production compared to WT (Figure 4). However, the *gonst1-1gonst2-1* double mutant had have a significant increase in both SA and H_2_O_2_, as compared to WT and *gonst1-1* alone (Figure 4).

**Figure 4:**
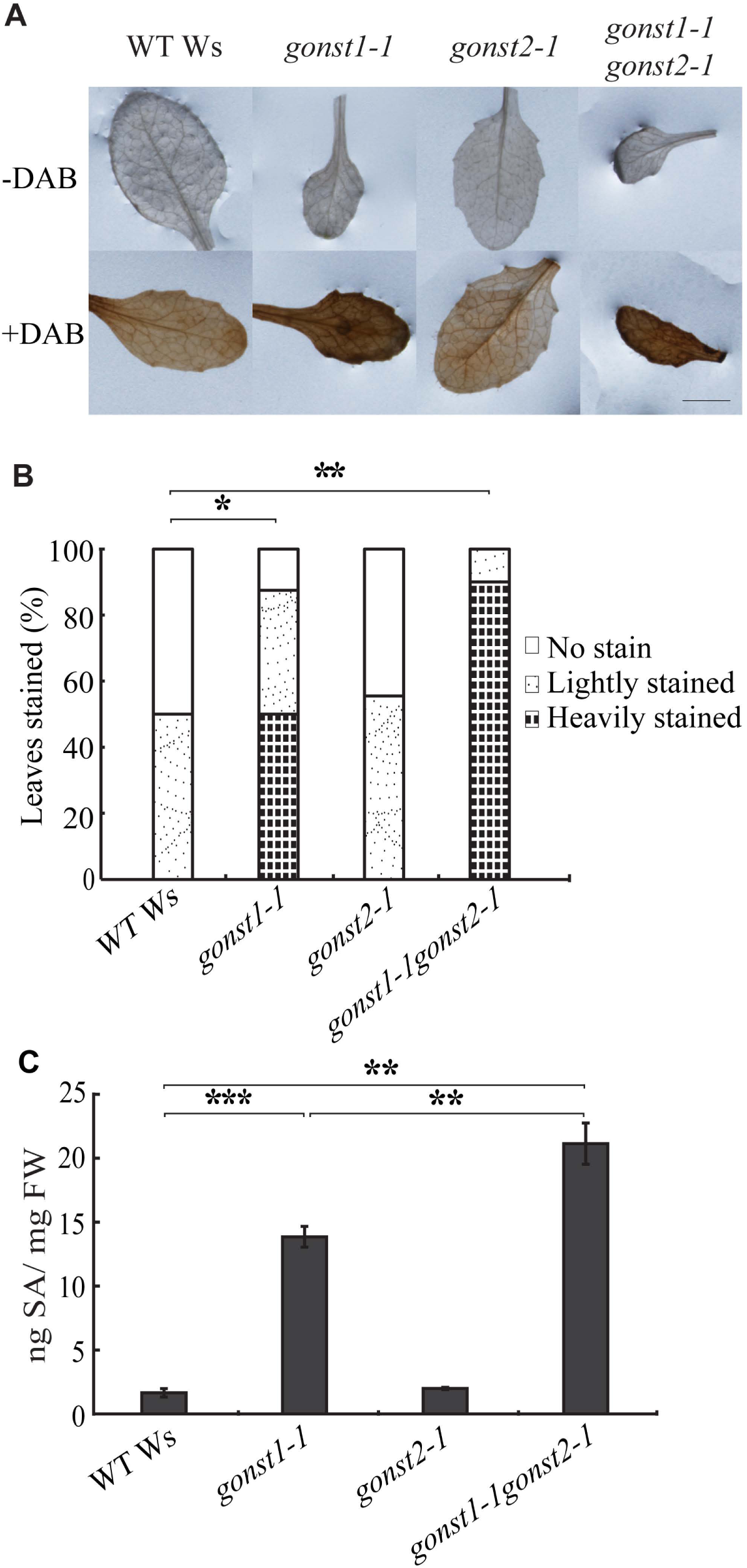
Loss of GONST2 enhances H_2_O_2_ and SA accumulation in the leaves of *gonst1* plants. (A) Intracellular H_2_O_2_ production was measured in 12 d old leaves using DAB staining, and (B) the leaves were scored for staining intensity. The Mann-Whitney test was used to determine significant differences. (C) Total SA in 12 d old leaves was quantitated by HPLC. Student’s *t*-test was used to determine significant differences. All data is mean ± SD of 3 independently grown biological replicates; asterisk indicate a significant difference between the two indicated genotypes. * *p* < 0.05, ** *p* < 0.01, *** *p* < 0.001.

### *gonst2* has increased resistance to biotrophic but not to necrotrophic pathogens

Plant pathogens can be divided into two major groups depending on their lifestyle strategies: necrotrophy, and biotrophy. Necrotrophic pathogens kill host cells and extract nutrition from the dead host, while biotrophic pathogens colonize living cells and obtain nutrition from living hosts (Hammond-Kosack and Jones, 1997). Classically, SA signaling triggers resistance against biotrophic pathogens, whereas a combination of jasmonic acid (JA) and ethylene (ET) signaling activates resistance against necrotrophic pathogens and these two pathways are mostly antagonistic (Robert-Seilaniantz et al., 2011). We wanted to test whether *gonst1* or *gonst2* plants show increased pathogen resistance, and whether this was generic, or specific to biotrophic pathogens. However, *gonst1* and *gonst1gonst2* rosette leaves are not suitable for pathogen assays, as they are fully senesced by ~20 days under normal conditions. Therefore, we tested *gonst2-1*, despite the lack of a significant increase in SA or ROS (Figure 4), since its rosette leaves are healthy. Indeed, *gonst2-1* showed significant increase of resistance against biotrophic *G. orontii* MGH (an Arabidopsis-adapted powdery mildew) (Figure 5A). However, there was no significant difference between WT and *gonst2-1* susceptibility to four different isolates of *Botrytis cinerea* (Figure 5B-E; Table 1).

**Figure 5:**
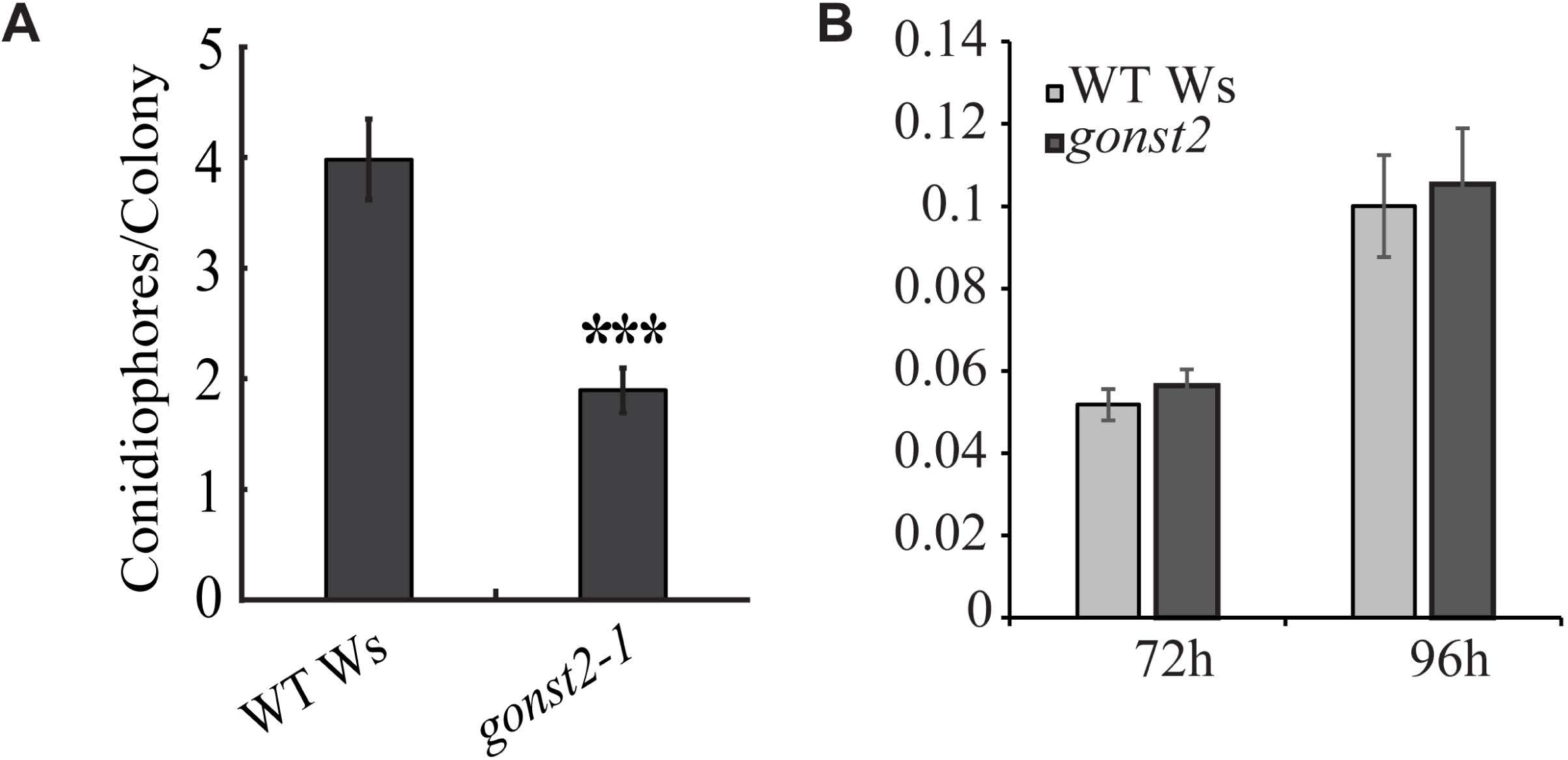
Susceptibility of *gonst2* plants to biotrophic and necrotrophic pathogens. (A) 5 days after inoculation with the biotroph *G. orontii*, leaves were harvested, stained with trypan blue, and conidiophores per colony counted. The data represents the mean of 12-30 leaves per genotype per experiment, scored in 3 independent experiments, ± SD. (Student’s *t*-test, * *p* < 0.05, ** *p* < 0.01, *** *p* < 0.001). ((B) 72 or 96 hours after inoculation with four phenotypically diverse *B. cinerea* isolates (1.02, 1.03, MEAP6G, and NobleRot), lesion size was measured. The data represent the mean of 105-166 leaves per plant genotype per experiment, + SE. No significant difference was detected between WT and *gonst2-*1 (F-test; Table 1).

### *gonst2* and *gonst1gonst2* have altered GIPC glycosylation

We have recently developed a simple thin layer chromatography (TLC) method which gives good separation of GIPCs which is primarily dependent on the nature and the degree of glycosylation (Murawska et al., in prep). Due to the small stature and tissue death in *gonst1-1* and *gonst1-1gonst2-1*, it was not possible to isolate GIPCs from whole plants. Therefore, we generated root-derived callus from all of the genotypes, isolated an enriched GIPC fraction, and performed TLC. The plates were stained with primuline and visualized under UV light (Figure 6A). As expected, *gonst1-1* shows a large shift in the mobility of the GIPCs, as compared to the Ws WT control, due to the loss of mannosylation as previously reported (Mortimer et al., 2013). A small fraction of the lower pool remains (marked with an arrow head). *gonst2-1* has a profile similar to WT with a small upper fraction of GIPCs (marked with an arrow head), and *gonst1-1gonst2-1* had essentially all GIPCs in the faster moving upper fraction.

**Figure 6:**
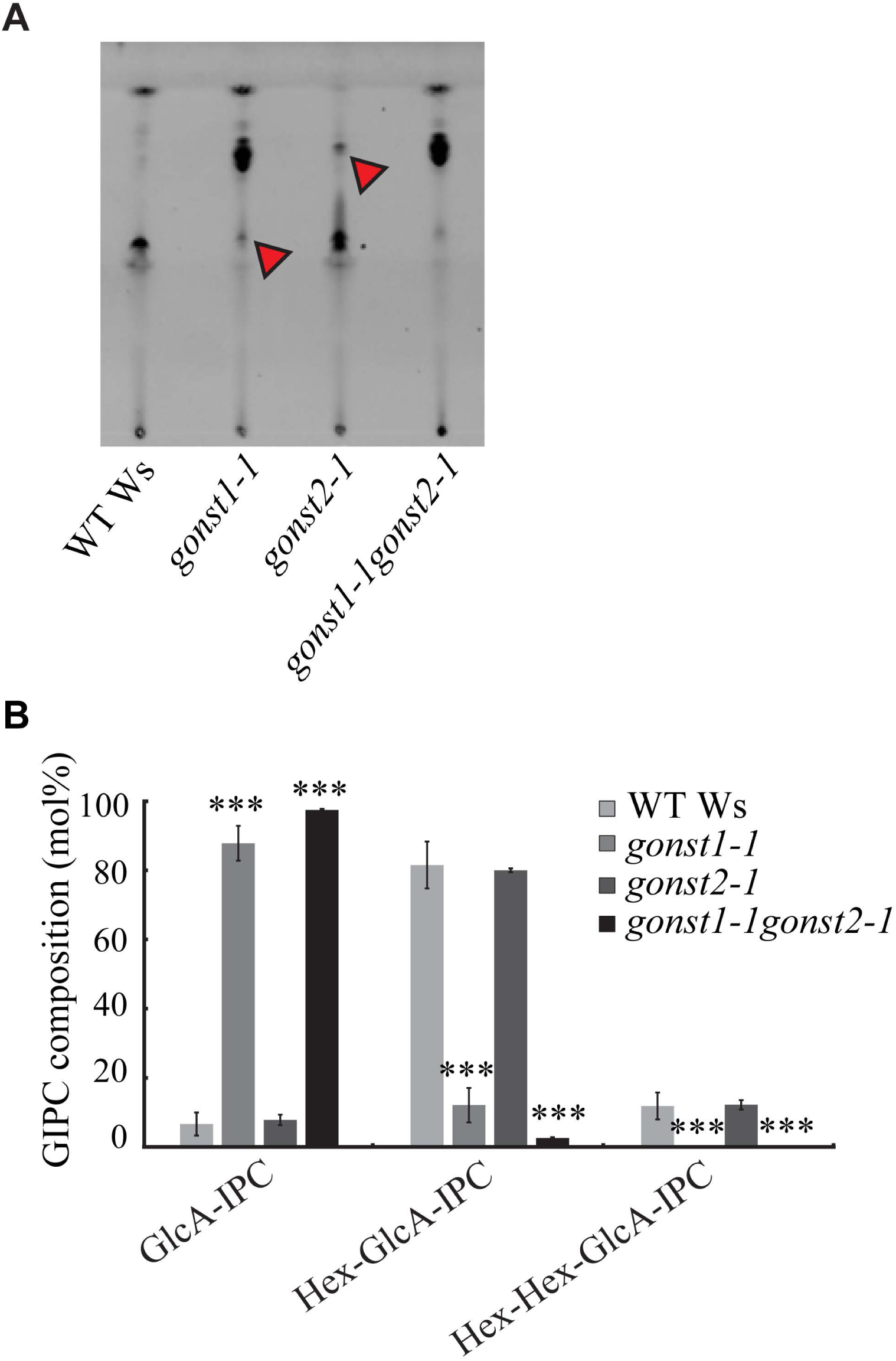
Glycan headgroup compositions of *gonst* GIPCs. (A) TLC of a GIPC-enriched membrane fraction which has been stained with primuline. Bands discussed in the text are marked with a red arrow head. (B) An enriched GIPC fraction was analyzed by LC-MS/MS MRM. The data here is collapsed to describe only the number of hexoses on the GIPC headgroup. All data is mean ± SD of 3 independently grown replicates of liquid grown cell culture; asterisk indicates significant difference from the WT (Student’s *t*-test, * p < 0.05, ** p < 0.01, *** p < 0.001). The full data set is shown in Supplemental Figure S4 and Supplemental Dataset S1.

To explore this further, we then used LC-MS/MS multiple reaction monitoring (MRM) to perform sphingolipidomics. No overall significant difference was detected in the glucosylceramides, hydroxyceramides or ceramides (Supplemental Figure S5, Supplemental Dataset 1). Plant GIPCs are enormously complex, due to the possible variations in fatty acid (FA), long chain base (LCB) and glycan structure. Since no changes in the lipid composition of the GIPCs were detected (Supplemental Figure S5), the GIPC data presented here has been simplified to just describe changes to the glycan headgroup Following the nomenclature described in (Fang et al., 2016)(Supplemental Figure S1), the data has been aggregated to show the relative amount of GIPCs containing either 0, 1 or 2 hexoses terminal to GlcA-IPC (Figure 6B). While *gonst2-1* did not show a significantly different GIPC headgroup profile from the WT, there was significantly less Hex_1_ GIPCs in *gonst1-1gonst2-1* than *gonst1-1*.

### *gonst2* and *gonst1gonst2* Golgi-synthesized cell wall polysaccharides are unaffected

Since GONST2 is a Golgi-localized nucleotide sugar transporter, we tested whether the loss of GONST2 could impact other glycosylation processes in the Golgi, other than GIPCs. The majority of cell wall polysaccharide biosynthesis, with the exception of cellulose and callose, occurs in the Golgi, so we first investigated the monosaccharide composition of the non-cellulosic polysaccharides of callus, leaves and stems by hydrolyzing an alcohol insoluble residue (AIR) cell wall preparation with trifluoroacetic acid (TFA) (Supplemental Figure S6). No significant difference was detected between *gonst2-1*, *gonst1-1gonst2-1* and the WT.

### *gonst2* and *gonst1gonst2* glucomannan structure and quantity is unchanged

Glucomannan is a Golgi-synthesized cell wall polysaccharide composed of β(1,4)-Man and −Glc, which is synthesized by the CSLA family of GTs. CSLA9 (the dominant mannan synthase in Arabidopsis vegetative tissue) requires GDP-Man and GDP-Glc for glucomannan synthesis (Dhugga et al., 2004; Liepman et al., 2005; Goubet et al., 2009), and it has been proposed that it has a luminal active site (Davis et al., 2010). However, no NST responsible for providing these substrates to the Golgi lumen has yet been identified. Loss of GONST1 does not affect glucomannan biosynthesis (Mortimer et al., 2013), GFT1 is a GDP-Fuc transporter (Rautengarten et al., 2016), and GGLT1 is a GDP-Gal transporter (Sechet et al., submitted). Since mannan is a relatively minor component of the cell wall (Handford et al., 2003), the monosaccharide analysis (Figure S6) may not reveal alterations to its quantity or the Glc:Man ratio. Therefore, we used Polysaccharide Analysis by Carbohydrate gel Electrophoresis (PACE) to investigate glucomannan quantity and structure (Handford et al., 2003). No difference was seen either in the type or quantity of the oligosaccharides released by hydrolysis of *gonst2*, *gonst1-1gonst2-1* and WT AIR by mannanases (Figure 7). Therefore, we can conclude that GONST1 and GONST2 are not providing substrate in the Golgi lumen for mannan biosynthesis.

**Figure 7:**
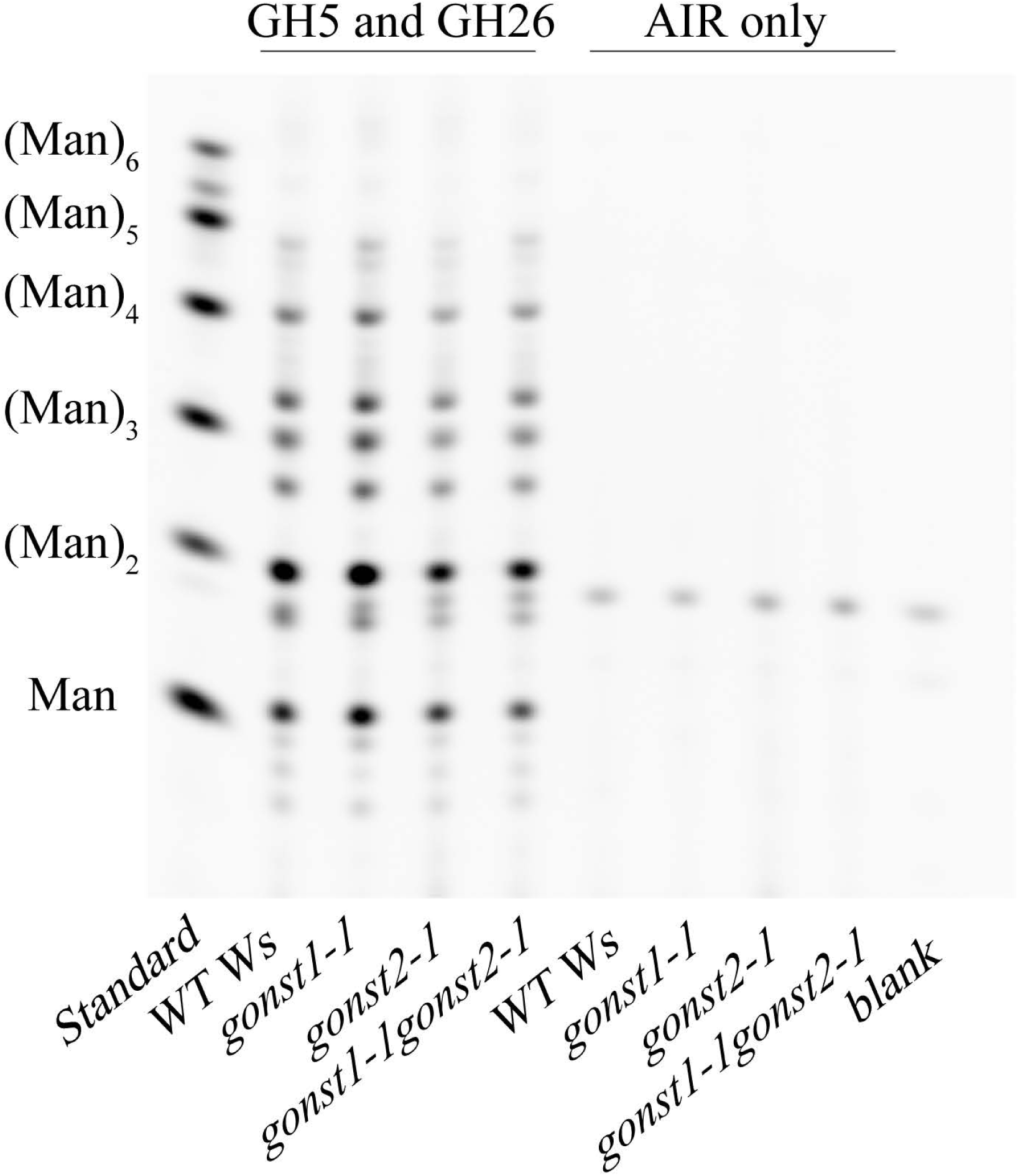
PACE fingerprint of mannan in stem cell walls of WT Ws, *gonst1*, *gonst2* and *gonst1gonst2*. Oligosaccharides released from AIR by mannanase digestion were derivatized with 8-aminonapthalene-1,3,6-trisulphonic acid (ANTS) and visualized by PACE. (Man)_1-6_ oligosaccharides were used as a standard. A representative gel from multiple experiments is shown.

### *gonst1gonst2* has less cellulose

Recently, we showed that a mutant in GIPC mannosylation (GMT1), has reduced cellulose (Fang et al., 2016). To test whether this phenotype is common to plants with altered GIPC mannosylation, we hydrolyzed the TFA-insoluble AIR fraction with sulfuric acid to release glucose derived from cellulose (Figure 8). *gonst1* had a significant decrease in upper and lower stem cellulose content, whereas callus and seedling were unaffected. *gonst2* did not show a significant difference in any tissue type analyzed compared to WT. However *gonst1gonst2* mutants showed a significant decrease in callus cellulose content (a tissue rich in primary cell wall), compared to the WT or single mutants. These data are consistent with a specific role for GIPC mannosylation in determining cell wall cellulose content.

**Figure 8:**
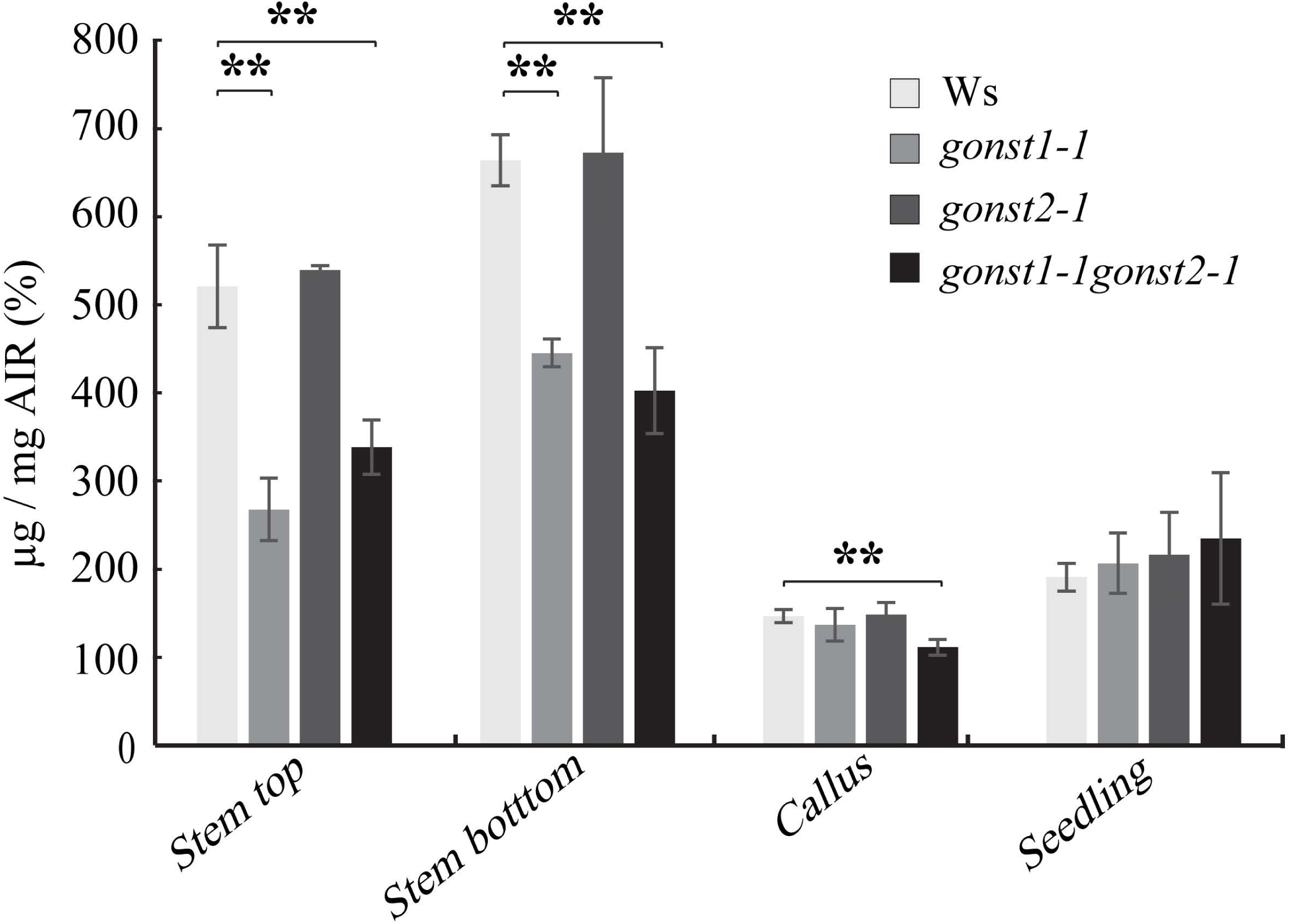
*gonst1gonst2* has reduced crystalline cellulose. Crystalline cellulose content was determined as the glucose content released by sulfuric acid treatment of the TFA-insoluble fraction of AIR. All data is mean ± SD of 3 biological replicates. Asterisk indicates significant difference from the WT (Student’s *t*-test, * p < 0.05, ** p < 0.01, *** p < 0.001).

### Expression of *GONST1pro:GONST2* in *gonst1* rescues growth and GIPC glycosylation

To test whether the phenotypic differences observed between *gonst1* and *gonst2* are due to functional differences or whether they are due to differences in expression level, we expressed *GONST2* CDS driven by either the *GONST1* promoter (*GONST1pro:GONST2*) or the *GONST2* promoter (*GONST2pro:GONST2*) in the *gonst1-1* background. Multiple independently transformed lines were selected for analysis. Analysis of T3 segregants revealed that some of homozygous *gonst1-1* plants had a restored growth phenotype. *GONST2* expression was analyzed by real-time RT-PCR (Figure 9A). The suppression of the *gonst1* growth phenotype was only apparent in those lines in which GONST2 expression was driven by the GONST1 promoter (Figure 9B). The rescue of the growth was reflected in the biochemical characterization of GIPC headgroup composition (Figure 9C). This result shows that GONST2 has the same function as GONST1, and that the function is cell type specific and/or dose dependent.

**Figure 9:**
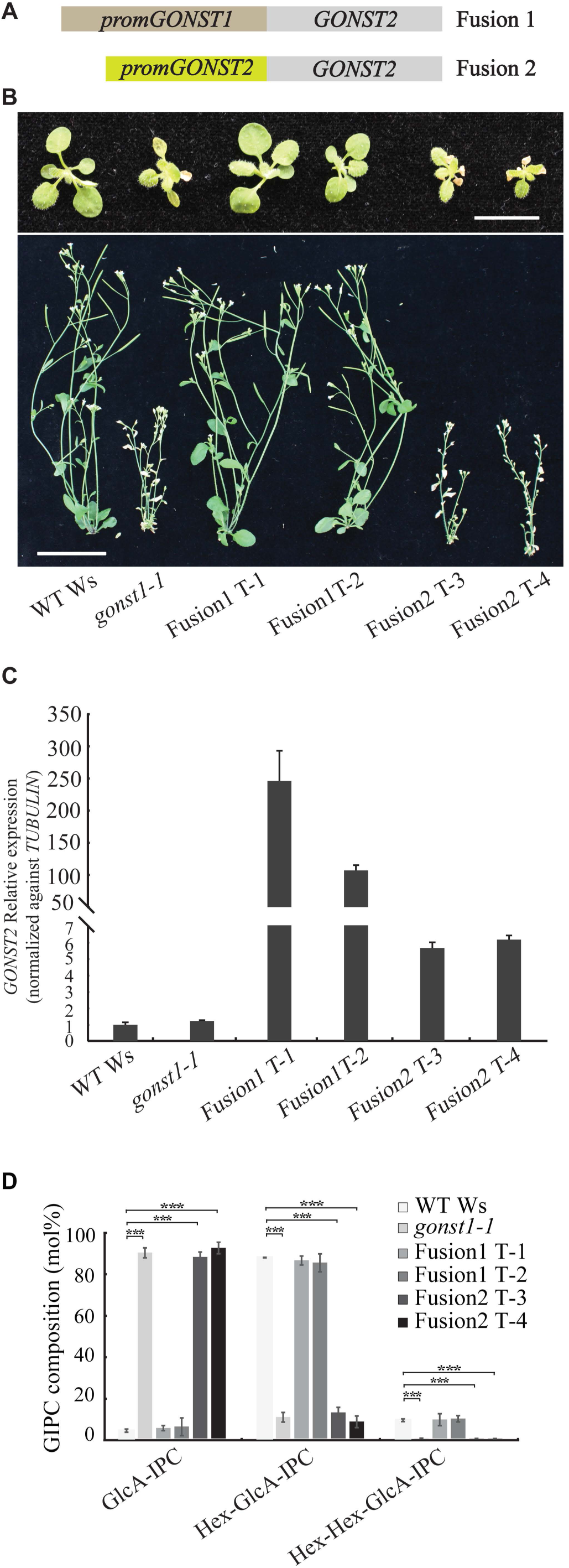
Expression of *GONST2* under the *GONST1* promoter in the *gonst1* background restores the dwarfed phenotype of *gonst1*. (A) Schematic of the two introduced constructs. (B) Top row: 15-day-old, agar grown WT, *gonst1-1*, Fusion1 T-1, Fusion1 T-2, Fusion2 T-3, and Fusion2 T-4 seedlings. Scale bar = 1 cm. Bottom row: 6-week-old WT, *gonst1-1*, Fusion1 T-1, Fusion1 T-2, Fusion1 T-3, and Fusion1 T-4. Plants were first grown on agar for 10 days, and then transplanted onto soil. Bar = 3 cm. (C) Gene expression analysis of *GONST2* relative to WT Ws and normalized against *TUBULIN* using Q-PCR. Values represent average of three biological replicates ± SD. (D) An enriched GIPC fraction was analyzed by LC-MS/MS MRM. The data here is collapsed to describe only the number of hexoses on the GIPC headgroup. All data is mean ± SD of 3 independently grown replicates of liquid grown cell culture. Asterisk indicates significant difference from the WT (Student’s *t*-test, * p < 0.05, ** p < 0.01, *** p < 0.001).

## Discussion

The aim of this research was to characterize the role of the final *bona fide* member of the GONST clade of four nucleotide sugar transporters, GONST2. We have now shown that, whilst *in vitro* it has been reported to transport all GDP-linked sugars (Rautengarten et al., 2016), *in planta* it has a specific role in providing GDP-Man for GIPC glycosylation. We also show that it is a functional homolog of GONST1. Since *gonst2* does not display the severe growth defects of *gonst1*, we were able to perform pathoassays to investigate the previously reported constitutive defense response of *gonst1* (Mortimer et al., 2013), and show that *gonst2* has increased resistance to the biotroph *G. orontii* but not the necrotroph *B. cinerea*.

Resistance to biotrophic pathogens, such as the powdery mildew-causing *G. orontii* is regulated by SA signaling (Wildermuth et al., 2001). Activation of SA signaling is often correlated with accumulation of reactive oxygen species including H_2_O_2_ (Herrera-Vasquez et al., 2015). On the other hand, resistance to necrotrophic pathogens such as *B. cinerea* requires JA/ET signaling, which mostly function antagonistically with SA (Robert-Seilaniantz et al., 2011). We found that the *gonst1gonst2* double mutant contains significantly increased SA and enhanced H_2_O_2_ accumulation. While uninfected *gonst2* did not show an increased SA level, it is possible that the SA level is enhanced in *gonst2* after *G. orontii* infection, contributing to the enhanced resistance to *G. orontii*. It has been reported that altered ceramide profiles are associated with altered phytohoromone levels, and thus with altered response to pathogens (Magnin-Robert et al., 2015). In this case, ceramide functions as a signaling component. While *gonst2* does not show a significant change to the ceramide pool, changes to the GIPC glycosylation may be enough to affect SA signaling and thus the response to *G. orontii*. Alternatively, a defect in membrane trafficking in *gonst2* may negatively impact *G. orntii* infection. *G. orontii* forms a specialized infection hypha called the haustorium in the host apoplast to establish infection. The haustorium is surrounded by host-derived membrane called the extrahaustorial membrane, which has modified endosomal characteristics (Inada et al., 2016). It has been shown that GIPCs are important for secretory sorting of proteins (Markham et al., 2011; Wattelet-Boyer et al., 2016), and therefore it may be that changes to a minor class of GPIC are enough to disrupt these processes, thereby negatively affecting *G. orontii* infection.

Cellulose content is decreased in *gonst1* and *gonst1gonst2* plants. Cellulose is synthesized at the plasma membrane by rosettes of CESA proteins which move through the plane of the plasma membrane (McFarlane et al., 2014). The rosettes are assembled in the Golgi and are delivered to the plasma membrane via the secretory system (Wightman and Turner, 2010). The reduced cellulose phenotype was also reported for *gmt1*, which has the same biochemical GIPC phenotype as *gonst1gonst2* (Fang et al., 2016). The reasons for this decrease are not clear. It is possible that the altered GIPC glycosylation affects trafficking of the rosettes to the plasma membrane, or alternatively, the change to plasma membrane composition affects CESA function. CESA proteins are S-acylated, and it has been suggested that this decoration may either localize proteins to lipid microdomains (which are rich in GIPCs) or even facilitate their formation (Konrad and Ott, 2015; Kumar et al., 2016). COBRA and COBRA-Like proteins which are also essential for normal cellulose biosynthesis are glycosylinositolphosphatidylinositol (GPI) anchored (Roudier et al., 2005). GPI anchored proteins are targeted to the outer leaflet of the plasma membrane, and to lipid microdomains. Therefore, correct GIPC glycosylation may be necessary for either CESA activity or localization and retention of GPI-anchored proteins in the plasma membrane.

It should be noted that alterations to cellulose content can affect susceptibility to some pathogens (Hernandez-Blanco et al., 2007; Malinovsky et al., 2014). For example, CESA3 mutants (a primary cell wall CESA) are more resistant to powdery mildews (Ellis and Turner, 2001; Cano-Delgado et al., 2003) and mutants in secondary cell wall CESAs are more resistant to necrotrophs (Hernandez-Blanco et al., 2007).

Mannan content is unchanged in the *gonst1gonst2* plants. It had been reported that CSLA9, unlike related GT2 proteins (CSLC4, CESAs) have a topology which results in a luminal active site (Davis et al., 2010). This would necessarily require a nucleotide transporter to provide GDP-sugars for mannan biosynthesis. However, none of the predicted GDP-sugar transporters seem to have this function *in planta* (Mortimer et al., 2013; Rautengarten et al., 2016)(Sechet et al., submitted). This implies that either the mannan synthases do not require a nucleotide sugar transporter, or that the transporter does not have a canonical GDP-binding motif.

Future work will include establishing how GIPC glycosylation affects these different membrane-based processes. For example, molecular dynamics could be applied to model the plant plasma membrane and understand how the GIPC glycan headgroup structure affect proteins movement within the membrane. It will also be interesting to understand what drives the differences in functionality of the NSTs in *in vitro* assays versus *in planta* function. Both GONST1 and GONST2 can transport all GDP-sugars when tested in liposome based assays (Mortimer et al., 2013; Rautengarten et al., 2016), but it is clear that they are highly specific *in vivo*. This could be mediated by substrate concentration, interaction with non-catalytic proteins, or interactions with the GT that utilizes the substrate (in this case GMT1 (Fang et al., 2016)). The recent crystal structure of the yeast Vrg4 NST provided new insights into how NST function is regulated (Parker and Newstead, 2017). To our knowledge, no plant NSTs have yet been structurally characterized, but we expect that this information will be critical for understanding NST specificity.

## Materials and Methods

### Materials

All constructs described in this publication are available upon request from the Joint BioEnergy Institute (JBEI)’s Inventory of Composable Elements (ICE) (registry.jbei.org). All chemicals are from Sigma Aldrich, unless otherwise noted. The GONST2 C-terminal YFP construct pEarleygate101 GONST2 was a generous gift from Dr. Carsten Rautengarten, Lawrence Berkeley National Laboratory under the 35S promoter, and was described previously (Rautengarten et al., 2016).

### Samples

All experiments were performed on at least three independently grown biological replicates unless otherwise stated.

### Phylogenetics

Protein sequences of Arabidopsis GONST2 were downloaded from TAIR (www.arabidopsis.org) and used in a BLASTp search (standard parameters) in NCBI. Species included were as follows: *Brachypodium distachyon*, *Zea mays*, *Oryza sativa*, *Camelina sativa*, *Populus euphratica*, *Eucalyptus grandis*, *Solanum lycopersicum*, *Nicotiana sylvestris* and *Glycine max*. Sequences were aligned with Clustal Omega (www.ebi.org) (standard parameters). The Phylip program set (v3.95) was used to build the tree, using standard parameters except where stated, as follows: seqboot (2000 replicates), proml (not rough analysis), consense and drawgram. All bootstrap probabilities were 1.0 with 2000 replicates.

### Plant material and growth conditions

The T-DNA line *gonst2-1* (FLAG_406C01; ecotype Ws; insertion into AT1G07290), as well as *gonst1-1* (FLAG_164D07; insertion into AT2G13650) were previously described in (Mortimer et al., 2013). A second independent null *GONST2* T-DNA insertion was not available, so two additional *gonst2* alleles were generated using CRISPR/Cas9 gene editing technology as described below. Arabidopsis seeds were surface sterilized and sown on solid medium containing 0.5x Murashige and Skoog salts including vitamins and 1% (w/v) sucrose. Following stratification (48 h, 4°C, in the dark), plates were transferred to a growth room (22°C, 100-200 µmol m^−2^ s^−1^, 14 h light/10 h dark, 60% humidity). After 2 to 3 weeks, plants were transferred to soil or Magenta boxes under the same conditions. For *G. orontii* experiments, plants were grown under a 12 h light/ 12 h dark photoperiod for 4-5 weeks before inoculation. Liquid callus cultures were derived from Arabidopsis roots and maintained as described previously (Prime et al., 2000).

### CRISPR Construct Building

Specific guide RNAs (gRNAs) for targeting *GONST2* gene were identified by searching CRISPR-PLANT (https://www.genome.arizona.edu/crispr/about.html) website (Xie et al., 2014) with the gene ID At1g07290. From the list of Class 0.0 and Class 1.0 gRNA candidates, GONST2_gRNA1 (5’ GAGATAACAGGCGTGACCAC 3’) and GONST2_gRNA2 (5’ GTTTGGTGGGTTCATTAAACA 3’) were further selected by manual alignment to ensure specific targeting of GONST2 but not the other GONST family members. Building of the CAS9-gRNA expression construct involved two steps. First, the entry vector containing pU6::GONST2_gRNA::scaffold::tNOS was built by PCR reactions with primers incorporating a specific gRNA sequence. PCR template and primers are listed in Table 1. The resulting three DNA fragments (i.e. gRNA Specific Piece1, gRNA Specific Piece2, Universal Piece 3) were assembled into the Entry vector by an In-Fusion reaction (In-Fusion HD, Clontech). Second, an LR recombination reaction (Gateway Cloning, Invitrogen) was performed to sub-clone pU6::GONST2_gRNA::scaffold::tNOS from the Entry vector into the expression vector C382, which contains the CAS9 expression cassette.

**Table 1:** ANOVA results for the various factors in the *Botrytis cinerea* infection experiment on WT and *gonst2* plants at 72 hours post inoculation. df = degrees of freedoms, SS = Type III Sums-of-Squares and *P* = estimated *P*-value.

| Sources of Variation | df | SS | *P* |
| --- | --- | --- | --- |
| Botrytis Isolates | 3 | 1.623 | <0.001 |
| WT vs *gost2* | 1 | 0.00963 | 0.3784 |
| Time | 1 | 0.04897 | 0.1171 |
| Isolate x Plant Genotype | 3 | 0.2676 | 0.8487 |
| Random Effects | df | LRT | *P* |
| Tray / Flat | 1 | 1.3844 | 0.2394 |
| Tray | 1 | 9.1055 | 0.002548 |

### Arabidopsis Transformation and Genotyping to Identify *gonst2-2* and *gonst2-3*

Agrobacterium strain GV3101 harboring C384 or C385 vectors was used to transform Arabidopsis plants with the floral dip method. T1 and T2 transgenic plants were selected on solid Murashige and Skoog medium supplemented with 1% sucrose and 50 μg/mL kanamycin. Two homozygous allelic mutant lines of *GONST2*, namely *gonst2-2* and *gonst2-3*, were identified at the T2 transgenic generation. Briefly, pooled leaf samples were harvested at 20 days post-germination. Genomic DNA was prepared with CTAB method (Doyle, 1987). DNA (100 ng) from each sample was used as the template for amplification of the target region using Phusion^®^ High-Fidelity DNA Polymerase (New England Biolabs, Ipswich, MA, USA). Primers are listed in Table 1. The amplicon was column purified before submitted for Sanger sequencing. Genotyping showed that *gonst2-2* line contained a single Thymine insertion at position 324; and *gonst2-3* line contained a single Adenine insertion at site 1889 of *GONST2* genomic sequence (start codon as position 1).

### Subcellular Localization

*Agrobacterium tumefaciens* (GV3101) transformed with either *35Spro*:*GONST2-YFP* or the Golgi marker Man49-GFP (Nelson et al., 2007) were co-infiltrated into 4 week old tobacco leaves. An additional *A. tumefaciens* strain carrying the p19 plasmid was also co-infiltrated to stabilize the transgene expression. Forty-eight hours after infiltration, the epidermal cells were removed from the tobacco leaves, fixed with formaldehyde and imaged using a Zeiss LSM 710 (Carl Zeiss, http://www.zeiss.com/) as previously outlined (Parsons et al., 2012). Image analysis and processing (scale bar, brightness, and contrast) were performed using IMAGEJ (Version 1.6r) (Schneider et al., 2012).

### Histochemical detection of H_2_O_2_

Detection of H_2_O_2_ was by endogenous peroxidase-dependent histochemical staining using 3,3-diaminobenzidine (DAB) as described in (Mortimer et al., 2013). Leaves of 15-day-old agar grown plants were submerged in 1 mL buffer (100 mM HEPES-KOH, pH 6.8) or 1 mg mL^−1^ DAB in buffer. After 4 min of vacuum infiltration, leaves were incubated at 22°C under a light intensity of 100-200 µmol m^−2^ s^−1^ for 6 h. Leaves were cleared for 30 min in 96% (v/v) ethanol solution at 70°C, and examined using a light microscope.

### Quantitation of salicylic acid (SA)

For total SA determination, 500 mg leaves were frozen and ground in liquid nitrogen. The powder obtained was mixed with 1 mL 80% (v/v) methanol and incubated for 15 min at 70°C. This step was repeated four times. Pooled extracts were centrifuged and filtered through Amicon Ultra centrifugal filters (10,000 Da MW cutoff, EMD Millipore, Billerica, MA). The conjugated SA in the filtered extracts was dried and the hydrolyzed in 1 N HCl at 95 °C for 3 h. The mixture was subjected to three ethyl acetate partitioning steps. Ethyl acetate fractions were pooled, dried *in vacuo*, and resuspended in 50% (v/v) methanol. SA was quantified using HPLC-electrospray ionization (ESI)-time-of-flight (TOF) MS. Details of the running condition were described previously (Eudes et al., 2013).

### *G. orontii* infection assay

*G. orontii* MGH was maintained on *pad4* leaves (Inada et al., 2016), and WT or *gonst2-1* leaves were inoculated using a settling tower method, as previously described (Plotnikova et al., 1998). Five days after inoculation, leaves were collected ad cleared in 99% ethanol, stained with Trypan Blue (250 µg mL-1 Trypan Blue in 1:1:1 glycerol:lactic acid:water) for ~15 minutes at room temperature, destained (1:1:1 glycerol:lactic acid:water) and visualized under a light microscope. Three independent inoculation experiments were performed, and 12-30 leaves were used to count numbers of conidiophores per colony, for each genotype, in each experiment.

### *B. cinerea* infection assay

*B. cinerea* inoculation followed a previously described protocol (Denby et al., 2004; Kliebenstein et al., 2005). We grew seedlings in a randomized complete block design on soil (SunGro Horticulture, Agawam, MA) growth chambers in 20°C, short-day (8h photoperiod) conditions. Spores were collected from mature *B. cinerea* cultures grown on canned peach plates and diluted to 10 spores/ µL in filter-sterilized 50% organic grape juice. At 7 weeks of age, we detached leaves from plants and arrayed them on 1% phytoagar by their order in the planting flats. We inoculated 4 uL spore solution droplets onto each leaf in a randomized complete block design, then incubated under 12 hour light / 12 hour dark at room temperature for 96 hours. The spore solution was continuously agitated to ensure equal distribution of spores. Digital images were taken at 48, 72, and 96 hours post inoculation. Lesion area was measured using custom R scripts along with the EBImage and CRImage packages (RDevelopment CORE TEAM, 2008; Pau et al., 2010; Failmezger et al., 2012).

### GIPC analysis by TLC

GIPCs were extracted as described in (Murawaska et al., in prep). Briefly, 200 mg of powdered lyophilized liquid-grown callus was added to 5 mL of the lower layer of isopropanol:hexane:water (55:20:25) and incubated at 50 °C for 15 min. Following centrifugation (500 × *g*, 10 min), the supernatant was transferred to a fresh tube, and the pellet was re-extracted with a further 5 mL of the lower layer of isopropanol:hexane:water (55:20:25). The supernatants were combined, dried under N2 and de-esterified by incubation with 33% (v/v) methylamine in ethanol:water (7:3) at 50 °C for 1 h. After centrifugation (500 × *g*, 10 min), the supernatant was retained, dried under N2 and incubated in 1 mL of chloroform:ethanol:ammonia:water (10:60:6:24) overnight at 21 °C with agitation. Samples were subjected to weak anion exchange chromatography as described in (Mortimer et al., 2013), and following elution from the cartridge were resuspended in chloroform:methanol:[4 M ammonium hydroxide in 1.8 M ammonium acetate] (9:7:2) and separated by thin layer chromatography (TLC) using high-performance-TLC Silica gel on glass plates (Merck) developed in the same buffer. GIPCs were visualized using primuline (Skipski, 1975).

### GIPC analysis by LC/MS

Total lipid for sphingolipidomics was prepared from lyophilized tissues (5-10 mg dry weight) using a methanol/butanol-based extraction coupled with weak alkaline hydrolysis and HCl treatment to remove glycerolipids and polysaccharides respectively, according to the previous report (Ishikawa et al., 2018). Each sphingolipid species was quantified using LC-MS/MS (LCMS-8030, Shimadzu, Kyoto, Japan) with the MRM mode targeting glucosylceramides, free ceramides and GIPCs with 0, 1 and 2 hexoses on GlcA-IPCs. The contents of Hex-GIPCs and ceramides were absolutely quantified by an internal standard-based calculation method, and GlcA-IPCs and Hex-Hex-GIPCs were relatively quantified using the calculation factors as Hex-GIPCs (Fang et al., 2016).

### Cell wall monosaccharide analysis

AIR was prepared according to (Mortimer et al., 2010) and 5 mg was hydrolyzed with fresh 2M trifluoroacetic acid (TFA; 400 µL, 1 h, 121 °C). The supernatant was removed, and the pellet washed twice with water (400 µL). The supernatant and washings were combined, dried in vacuo, and analyzed by HPAEC-PAD as previously described (Fang et al., 2016). The TFA-insoluble pellet was subjected to Saeman hydrolysis. Briefly, following incubation in 72 % (v/v) sulfuric acid (63 µL, 21 °C, 1 h), water was added to each sample to give a final sulfuric acid concentration of 1 M and incubated at 100 °C for 3 hours. The samples were then neutralized with barium carbonate, to precipitate the sulfate ions, and the Glc content measured by HPAEC-PAD as above.

### Mannan structural analysis using PACE

PACE was performed according to (Goubet et al., 2009) with slight modifications. Briefly, AIR (500 µg) was incubated with concentrated NH3 for 30 min at 21 °C, and then dried in vacuo. Following resuspension in ammonium acetate buffer (0.1 M, 500 µL, pH 6.0) samples were incubated for 14 h at 21 °C with an excess of the mannanases CjMan5A and CjMan26A (a kind gift from Professor Harry Gilbert, University of Newcastle, UK. The released oligosaccharides were derivatized with 8-aminonaphthalene-1,3,6-trisulfonic acid (Invitrogen) with 2-picoline-borane as the reducing agent, and separated by electrophoresis in large-format polyacrylamide gels. Gels were visualized using a Syngene G:BOX gel doc system (Synoptics, Cambridge, UK), equipped with long-wave UV transilluminator bulbs and appropriate filters.

### Promoter swap

The GONST1 promoter (1.3kb upstream of the start codon) and GONST2 promoter (1.0kb upstream of the start codon) were was amplified by PCR from Col-0 genomic DNA, and cloned into the binary vector pCAMBIA1305 to obtain pCAMBIA1305 *GONST1pro* and pCAMBIA1305 *GONST2pro*. Full-length cDNA of *GONST2* were amplified by PCR and cloned into pCAMBIA1305 *GONST1pro* and pCAMBIA1305 *GONST2pro* to obtain pCAMBIA1305 *GONST1pro:GONST2* (Fusion 1) and pCAMBIA1305 *GONST2pro:GONST2* (Fusion 2). Constructs were transformed into Agrobacterium as above and used to transform homozygous *gonst1-1* plants. T3 plants which were confirmed to be homozygous for the *gonst1-1* T-DNA insertion ((Mortimer et al., 2013) were analyzed.

## Accession numbers

GONST1 (Q941R4.2) and GONST2 (AEE28103.2).

## Supplemental Material

Supplemental Figure S1: Example GIPC structure, including the nomenclature used in this manuscript

Supplemental Figure S2: Phylogenetic characterization of Arabidopsis GONST family.

Supplemental Figure S3: Tissue-specific expression of *GONST2* and *GONST1*.

Supplemental Figure S4: Characterization of *gonst2-2* and *gonst2-3.*

Supplemental Figure S5: Sphingolipidomic analysis of WT, *gonst1-1*, *gonst2-1* and *gonst1-1gonst2-1* callus.

Supplemental Figure S6: Monosaccharide composition of non-cellulosic cell wall polysaccharides.

Supplemental Dataset S1: Sphingolipidomic data.

Supplemental Dataset S2: HPAEC-PAD data.

Supplemental Table S1: Oligonucleotide primers used in this project.

## Acknowledgements

The authors would like to thank Professor Taku Demura, Dr Misato Ohtani and the rest of the Demura research team for their support at RIKEN.

## One Sentence Summary

Loss of GONST2, a Golgi nucleotide sugar transporter, alters sphingolipid glycosylation and increases resistance to biotrophic, but not necrotrophic, pathogens.

## Author Contributions

JCM and BJ designed the research. BJ, JCM, FA, GM and RP carried out the research. TI performed sphingolidomics; NS and NI performed pathogen assays; EB analyzed salicylic acid quantities; YL generated CRISPR lines; XY analyzed mannan structure by PACE. JCM, BJ, TI, NS, NI, MKY, DL, and PD analyzed data.

